# The Geometry of Masking in Neural Populations

**DOI:** 10.1101/665174

**Authors:** Dario L. Ringach

## Abstract

We introduce a geometric approach to study the representation of orientation by populations of neurons in primary visual cortex in the presence and absence of an additive mask. Despite heterogeneous effects at the single cell level, a simple geometric model explains how population responses are transformed by the mask and reveals how changes in discriminability and bias relate to each other. We propose that studying the geometry of neural populations can yield insights into the role of contextual modulation in the processing of sensory signals.

## Introduction

Individual neurons in primary visual cortex respond to stimulation within restricted areas of the visual field, which define their classical receptive fields^1–3^. These responses can be modulated by contextual stimuli presented within the classical receptive field or in the surrounding regions^4–6^. Cross-orientation and surround suppression are two well-known examples of contextual modulation^5,7–21^.

The role that contextual modulation plays in cortical function remains an open question. Some have considered such interactions to be directly involved in image processing, such as the detection and enhancement of smooth, spatially extended contours^22–37^. Others maintain that the fundamental goal of contextual modulation is to generate a sparse, efficient representation of natural images^6,38–45^. Distinguishing between these theories is not straightforward, as the their goals are not mutually exclusive^6^.

Here we focus on how contextual modulation transforms the activity of neural populations. Contextual modulation has been studied extensively in single neurons, leading to the development of the influential normalization model^6,46,47^. We have recently shown, however, that key properties derived from the classic formulation of normalization, such as contrast and subspace invariance, do not strictly hold at a population level^48^. Thus, we and others^49^ see new opportunities in the study of contextual modulation at the level of neural populations. The present study follows up on this line of work by studying how the coding of orientation by neural populations is transformed in the presence of a mask.

## Results

### Measurement of population responses in masked and unmasked conditions

We measured the responses of neural populations in mouse primary visual cortex using two-photon imaging (**Methods**). Mice were head-restrained but otherwise free to walk on a rotating wheel. The visual stimulus consisted of two conditions (**Fig 1A**). In the *unmasked condition*, a full-field sinusoidal grating was presented while its orientation changed linearly with time *θ* = *πt*/*T* with a period *T* = 10s. This type of continuously rotating stimulus has previously been used to measure orientation maps^50^. In the *masked condition,* the same rotating stimulus was presented superimposed on top of a mask consisting of a sinusoidal grating oriented vertically. We estimated the spiking responses of neurons using a standard processing pipeline involving image registration, signal extraction and deconvolution^51^. The periodic nature of the stimulus was evident in the temporal responses of cells (**Fig 1B**). This is because neurons tuned to one orientation respond once per cycle. As described in earlier studies^52^, locomotion modulated the overall responses of the population (**Fig 1B**, shaded regions).

**Fig 1.**
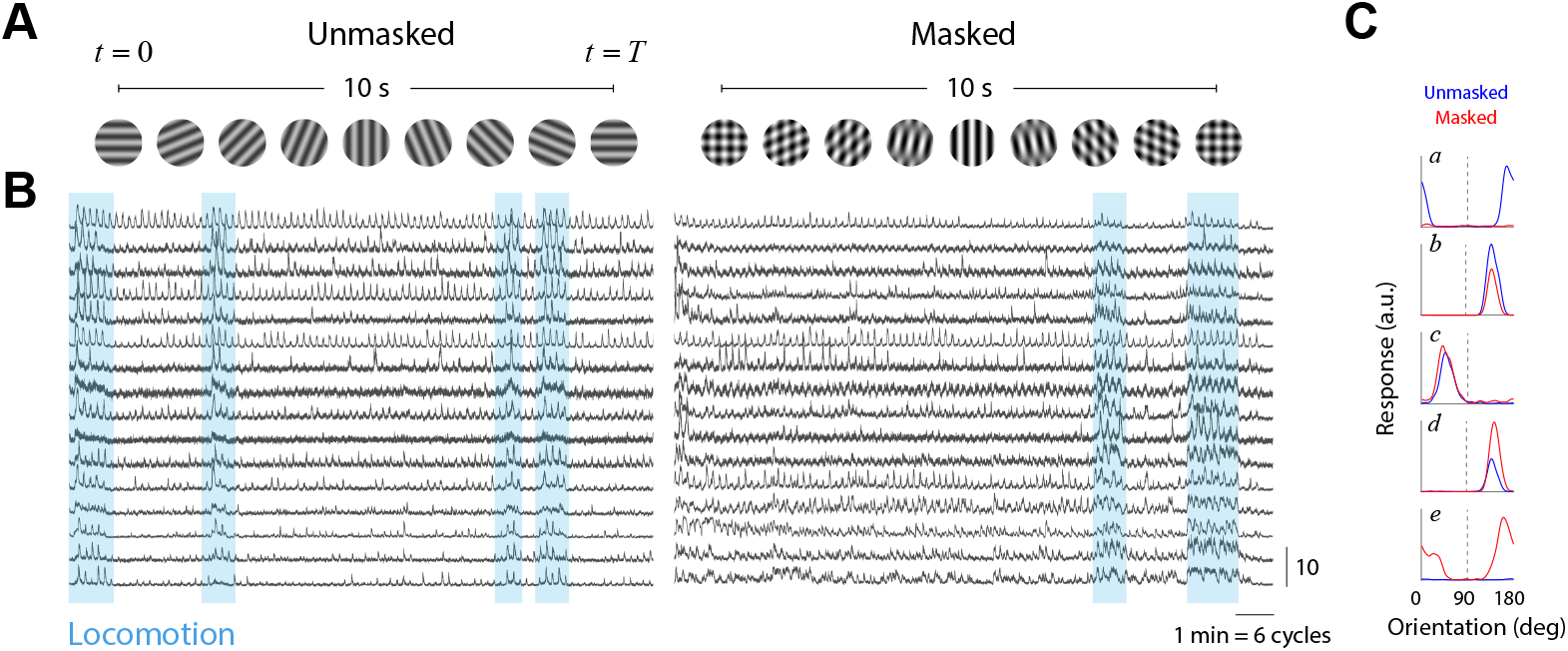
Measurement of population responses in masked and unmasked conditions. (**A**) Structure of the visual stimulus. Each of the lines show a single period of the stimulus in unmasked and masked conditions. (**B**) Samples of responses by individual neurons in both conditions. Some cells responded very well in the unmasked conditions (top traces) while others gave a weak response (bottom traces). Periods of locomotion enhanced the overall responsivity of the population (shaded regions). Traces are plotted on a z-scored scale (vertical bar = 10). Horizontal bar represents 1min of stimulation (or 6 periods of the orientation cycle). (**C**) Example of cell responses in unmasked and masked conditions. Each trace shows the response of a neuron over the stimulation cycle after correction for neural delay, so they can be interpreted as a sweep of the orientation tuning curve of the neuron. The dashed line indicates the orientation of the mask. Blue traces represent the responses in the unmasked condition, while red traces represent responses in the masked condition.

### Single neuron responses in masked and unmasked conditions are heterogenous

We computed the average response of neurons in the unmasked and masked conditions over the cycle of the stimulus (**Fig 1C**). The temporal responses were shifted by the mean stimulus-response delay (see **Methods**). After this correction, the temporal profile of the response can be interpreted as an estimate of the tuning curve of the neuron. In this representation, the mask is present at an orientation of 90 deg (**Fig 1C**, dashed lines; subsequent figures omit the location of the mask to avoid clutter).

We observed heterogeneity of responses at the single cell level. Some cells were well tuned to orientation in the unmasked condition but were completely suppressed by the addition of the mask (**Fig 1C*a***). Others did show such dramatic suppression, but responded with a scaled down version of their unmasked responses (**Fig 1C*b***) – a behavior consistent with the normalization model^6,46,53^. Some cells showed little or no difference between the responses in the two conditions (**Fig 1C*c***). Another group saw their unmasked responses scale up by the mask (**Fig 1C*d***). Finally, somewhat surprisingly, a set of neurons showed very weak responses in the unmasked condition but responded vigorously in the presence of the mask (**Fig 1C*e***)^48^.

We studied the range of behaviors in single cells (**Fig 1C**) by comparing the mean response the of *i* — *th* neuron over the stimulation cycle between unmasked and masked conditions, which we denote by 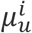 and 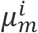 respectively (**Fig 2A**). We found a significant anti-correlation: the stronger a neuron responded in the unmasked condition the weaker its response was in the masked condition and vice-versa (*n* = 3920, *r* = −0.55, *p* = 5.6 × 10^−312^). We refer to neurons at the extremes of the distribution of behavior as *grating* and *plaid* cells (**Fig 2A**, shaded areas). These groups were formally defined as the cells attaining the 10% lowest (grating cells) and highest (plaid cells) ratios of 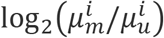 (**Fig 2A**, inset). We note these groups represent behaviors found at the extremes of a unimodal distribution (there is no evidence of discrete classes of neurons). Grating and plaid cells had different preferred orientations. Grating cells were preferentially tuned to the orientation orthogonal to the mask in both conditions (**Fig 2B,** left panels). In grating cells, the introduction of the mask scaled down the responses by about a third but did not affect tuning (note the different *y*-scales in **Fig 2B**). This is the type of responses one might expect from the classic normalization model^46,53^. Plaids cells, on the other hand, where preferentially tuned to the orientation of the mask (90 deg) when probed with grating stimuli in the unmasked condition, although their responses were relatively weak (**Fig 2B**, top right). Instead, and somewhat surprisingly, these cells were most responsive to the orthogonal orientation (0 deg) under the presence of the mask (**Fig 2B,** bottom right) – in other words, they responded best when the stimulus was a plaid with orthogonal components.

**Fig 2.**
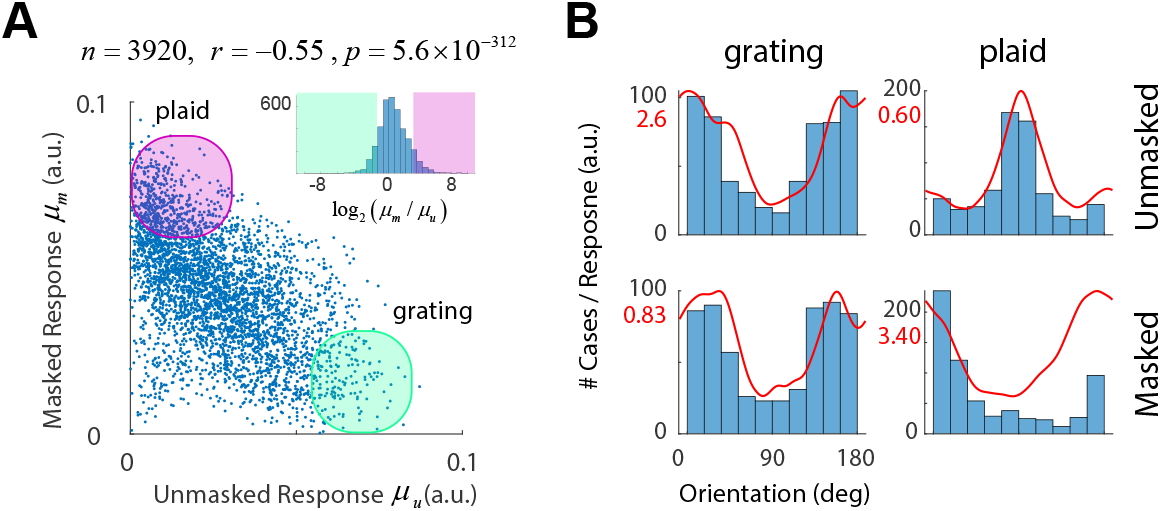
Characterization of responses in single neurons. (**A**) Anti-correlation between responses of neurons in masked and unmasked conditions. The mean responses of cells in the unmasked condition, *μ*_*u*_, are anti-correlated with the responses in the masked condition, *μ*_*m*_. The inset shows the distribution of log_2_(*μ*_*m*_/*μ*_*u*_). Cells at the extreme of this distribution are termed grating (shaded green) and plaid (shaded pink) neurons. (**B**) Average tuning of grating and plaid cells in unmasked and masked conditions. The histograms show the distribution of the preferred orientation of the neurons in each case. The red traces show the average tuning of neurons in each condition. The y-axis is labeled by cell count (in black) or by the amplitude of the responses (in red).

### A geometric framework to study contextual modulation in neural populations

The data in unmasked and masked conditions can each be represented as a matrix where the columns represent the tuning function of each cell (**Fig 3A**). To ease visualization, we ordered neurons by their preferred orientation. The rows of the matrix represent the population response to a given orientation. We denote the mean population responses as a function of orientation in the unmasked and masked conditions by *r*_*u*_(*θ*) and *r*_*m*_(*θ*), respectively. These vectors can be thought to describe parametric (closed) curves in a high dimensional space as *θ* ∊ [0, *π*] traverses the orientation domain (the dimension being the number of neurons in the population). We aim to understand the shape of these curves, the nature of the transformation *T*: *r*_*u*_(*θ*) → *r*_*m*_(*θ*) introduced by the mask, and how the outcome affects the discriminability of stimuli and biases the estimation of orientation in the masked condition.

**Fig 3.**
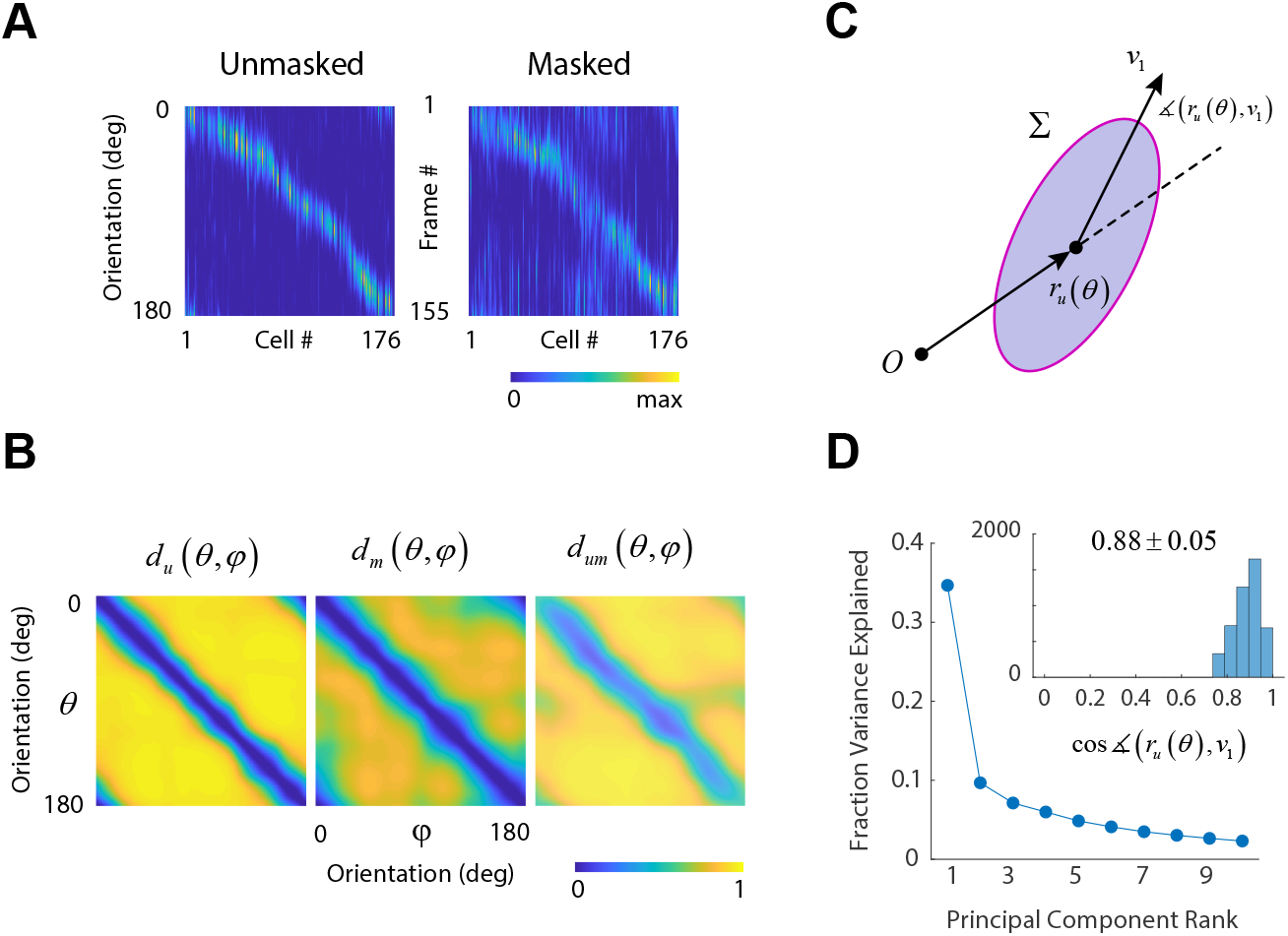
Characterization of population responses. (**A**) Responses of a population of neurons in the unmasked and masked conditions. Cells were ordered according to their preferred orientation, thus resulting in a diagonal structure. The rows for these matrices represent the population responses in the unmasked and masked conditions, *r*_*u*_(*θ*) and *r*_*m*_(*θ*). These curves describe a close curve as *θ* describes one cycle. (B). The intrinsic geometry of the curves is captured by the cosine distances between the representation of two orientations in the unmasked condition, *d*_*u*_(*θ*, *φ*) (left panel), and unmasked condition, *d*_*m*_(*θ*, *φ*) (middle panel). The relative positions of the curves with respect to each other is measured by the cosine distance between *r*_*u*_(*θ*) and *r*_*m*_(*φ*), denoted by *d*_*um*_(*θ*, *φ*) (right panel). (**C**). Schematic showing a response *r*_*u*_(*θ*) along with the covariance matrix and the direction of the first eigenvector, *v*_1_. (**D**) The first eigenvector/eigenvalue captured about a third of the total energy of the noise and the direction of the first eigenvector was very close to that of the response itself. The inset shows the distribution of the correlation coefficient between *r*_*u*_(*θ*) and *v*_1_ for all our experiments and orientations tested.

In what follows we denote by *d*_*u*_(*θ*, *φ*) the cosine distance between *r*_*u*_(*θ*) and *r*_*u*_(*φ*) (**Fig 3B,** left). The cosine distance is one minus the cosine of the angle between the two vectors. Because these vectors have positive entries representing a spike rate, the distance is bounded between zero and one. Similarly, we define *d*_*m*_(*θ*, *φ*) as the cosine distance between *r*_*m*_(*θ*) and *r*_*m*_(*φ*) (**Fig 3B**, middle). Under certain assumptions about the uniformity of the noise, the measurements *d*_*u*_(*θ*, *φ*) and *d*_*m*_(*θ*, *φ*) capture the ability of the population to discriminate between two angles in each condition. Finally, *d*_*um*_(*θ*, *φ*) denotes the cosine distance between the population representation of *θ* in the unmasked condition and the representation of *φ* in the masked condition. This measure captures the relative positions of the two curves and will induce biases in the estimation of orientation in the presence of a mask. Namely, non-zero biases result when the structure of *d*_*um*_(*θ*, *φ*) is not perfectly diagonal (**Fig 3B**, right). In the sequel, we also denote the normalized population vectors by 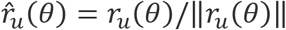 and 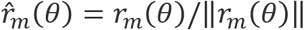.

### Orthogonality of signal and noise subspaces

We selected cosine distance as a metric because a substantial component of neural variability in the population response occurs along its direction^54^. To show this, we computed the mean and the covariance of the responses, *r*_*u*_(*θ*) and ∑_*u*_(*θ*). For each orientation, we compared the direction of the population response with the direction of the largest eigenvector, *v*_1_(*θ*), of the covariance matrix (**Fig 3C**). The largest eigenvector accounted for nearly a third of the variability (and more than three times the variance accounted for by the next largest eigenvalue) (**Fig 3D**) and its direction was very similar to that of the largest eigenvector – the correlation coefficient between *r*_*u*_(*θ*) and *v*_1_(*θ*) was 0.88 ± 0.05 (mean ± 1SD) (**Fig 3D,** inset). This large component of variability is due to fluctuations in behavioral state which modulates the gain of the response vector^52,55,56^. An implicit assumption behind the adoption of the cosine metric is that the orientation of the stimulus is coded by the *direction* of the population vector^46,57^. Thus, the result can be rephrased by stating that for each orientation, the direction of largest variability, 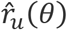, is orthogonal to the direction of the encoding, 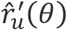, which is tangent to the unit sphere.

### Multidimensional scaling of population responses

To gain insight about the geometry of the curves and their relative positions we visualized them using multi-dimensional scaling using the cosine distance as a metric (**Fig 4**). The curves represent the embeddings of 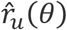 (blue) and 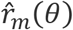 (red) in 3D space, while the spheres of matching colors indicate the point where the stimulus cycle begins. We define the mean population response over the entire stimulation cycle as the *white point*, which we denote by denote by *μ*_*u*_ and *μ*_*m*_. The grey arrows depict the shift of the white points between unmasked and masked conditions, with the stem of the arrow positioned at *μ*_*u*_ and the head at *μ*_*m*_. These examples are typical of what we observed in our experiments.

**Fig 4.**
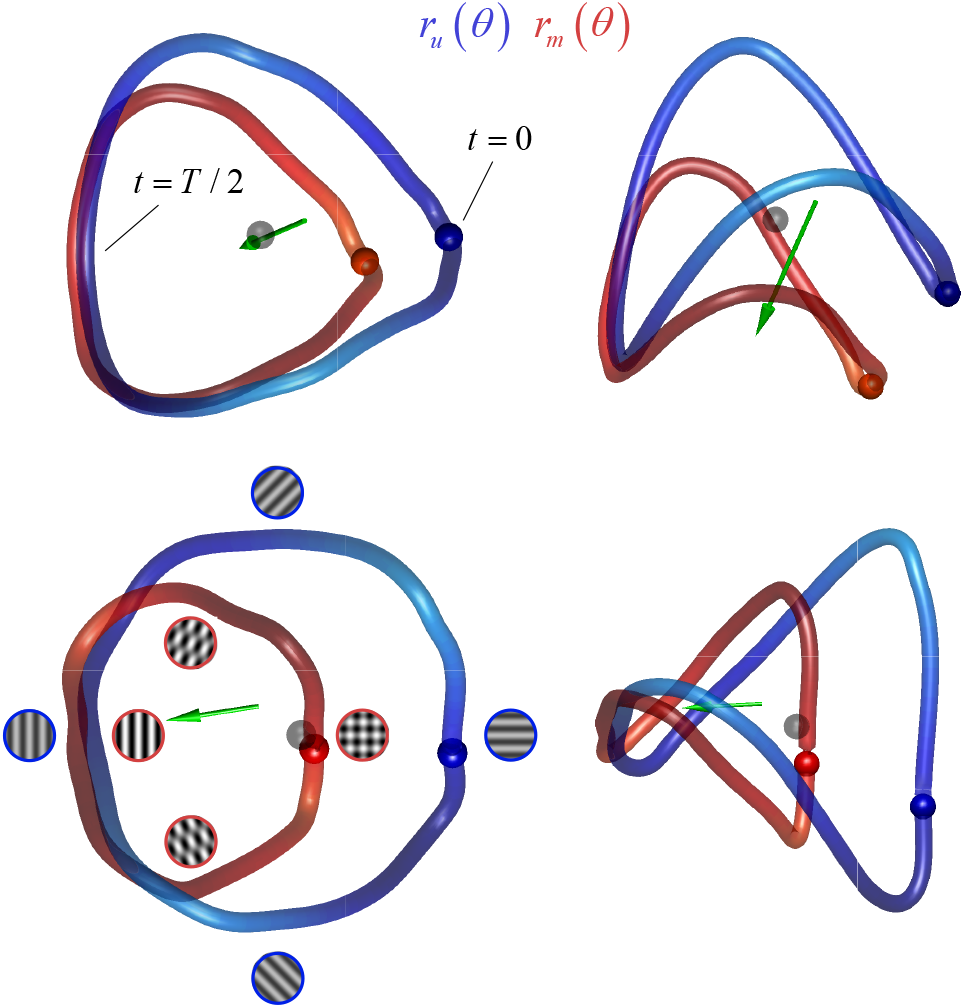
Multidimensional scaling of population responses in unmasked and masked conditions. Each row shows two viewpoints of the result of one experiment. The curves were obtained by doing multidimensional scaling simultaneously on the population responses in unmasked and masked conditions into 3D space using the cosine distance as a metric. The blue curve shows *r*_*u*_(*θ*) and the red curve shows *r*_*m*_(*θ*). The gray sphere represents the origin, and colored spheres represent the beginning of the cycle. The green arrows represent the shift in the white point between conditions. The stimuli represent the patterns at different locations on the curves for the two conditions (blue outline = unmasked condition; red outline = masked condition).

The shapes of 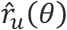 and 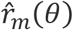 are similar, with the masked representation being a scaled down version of the original. The curves are farthest from each other at the beginning of the cycle, when the pattern in the masked condition consists of an orthogonal plaid and the one in the unmasked condition is a horizontal grating. The two curves are closest to each other near the middle of the cycle, when the pattern in the masked condition is a vertical grating with 100% contrast and the one in the unmasked condition is a vertical grating with 50% contrast. The curve 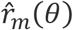 appears to be rotated away from that of 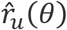, with the axis of rotation passing near the representation the mask. These features were consistent across our experiments suggesting that a scaling and rotation may explain the transformation of 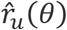 into 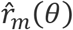 induced by the mask. Of course, these visualizations ought to be interpreted with caution, as they are only approximate representations of the geometry of high dimensional objects. Thus, we must check these first impressions of the geometry by doing appropriate calculations in the native space.

### Masking shrinks and rotates population responses

To verify our perception that curves are shrinking we computed their lengths^58^, 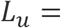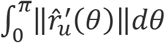 and 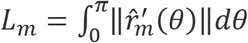. The arguments represent the angular velocity at which the population changes its orientation and represent a measure of discriminability between nearby angles. The length, therefore, represents local discriminability summed over all orientations^58,59^. The mask had the effect of reducing the overall length of the curves by a factor of 0.84 ± 0.05 (mean ± 1SD) (**Fig 5A**). As we will soon demonstrate, this shrinkage is not uniform, but peaks near the orientation of the mask.

**Fig 5.**
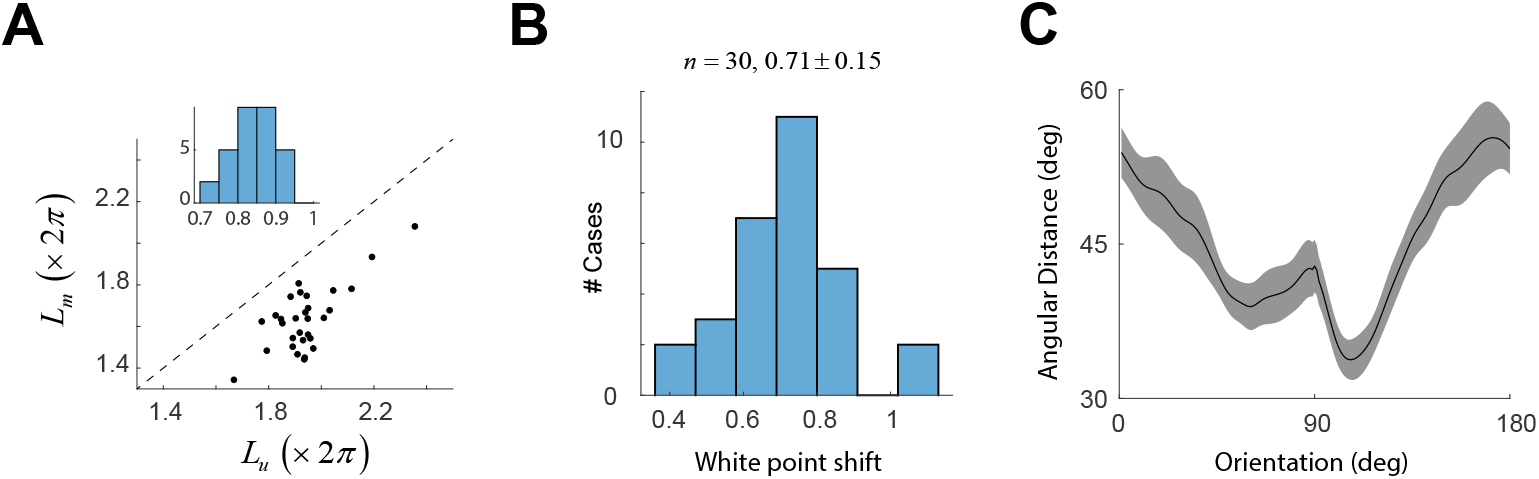
Basic geometric properties of population representations in unmasked and masked conditions. (**A**) Shrinkage of the length of the curves by the introduction of the mask. Scatterplot shows the lengths of the curves in unmasked (*L*_*u*_) and masked (*L*_*m*_) conditions. Dashed line represents the unity line. Inset shows the distribution of *L*_*m*_/*L*_*u*_ across all experiments. (**B**) Distribution of white-point shift (Δ) across all experiments. (**C**) Measurements of the angle between *r*_*m*_(*θ*) and the plane span {*r*_*u*_(*θ*), *r*_*u*_(*π*/2)} across all experiments. Solid line represents the mean, while the shaded area represents ± 2 SEM.

To verify our impression that the mask induces a change in the direction of the mean population activity, we defined the white-point shift as Δ= *d*(*μ*_*u*_, *μ*_*m*_)/((*ρ*_*u*_ + *ρ*_*m*_)/2). Here, *ρ*_*u*_ represents the average radius of the curve in the unmasked condition, calculated as 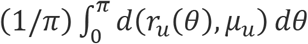, and a corresponding definition applies to *ρ*_*m*_. In other words, we measure the shift of the white point in terms of the average radius of the curves. Across the population we find Δ= 0.71 ± 0.15 (mean ± 1SD) – a relatively large fraction (**Fig 5B**), which is consistent with the visualizations from multi-dimensional scaling. We will see this shift is important because it is partly responsible for generating biases in the estimation of orientation in the masked condition.

### Rejection of the linear combination model

With the geometric formalism in place, we can test a common model of population responses, which postulates that the response to a plaid can be written as a linear combination of the population responses to the individual components^48,60^. The implication for our experiment is that *r*_*m*_(*θ*) ∊ span {*r*_*u*_(*θ*), *r*_*u*_(*π*/2)} (recall the mask has orientation *π*/2). One way to test the prediction is to measure the angle formed by the vector *r*_*m*_(*θ*) and the plane span {*r*_*u*_(*θ*), *r*_*u*_(*π*/2)}. The results show a significant departure from the prediction, with angular deviations larger than 30 deg and significantly higher than zero (*p* < 0.005, bootstrap, see **Methods**) (**Fig 5C**). Thus, the present data rule out the linear combination model, thereby confirming and extending a prior result^48^.

### The mask impairs discriminability and biases the decoding of orientation

Next, we analyzed changes in discriminability induced by the mask. Discriminability between any two orientations depends both on the distance between the mean population vectors and the statistics of the noise. As mentioned above, if the statistics of the noise are uniform in the sense that they translate with the direction of the population, we expect discriminability to be proportional to the distances *d*_*u*_(*θ*, *φ*) and *d*_*m*_(*θ*, *φ*). Nevertheless, given we have ~100 cycles we were able to compute a proper *d-prime* measure for both masked and unmasked conditions, which we denote by *D*_*u*_(*θ*, *φ*) and *D*_*m*_(*θ*, *φ*) (see **Methods**) (**Fig 6A**). To measure local discriminability (or just noticeable differences) we defined the threshold for detection in the unmasked condition *T*_*u*_(*θ*) as the minimal angle Δ such that *D*_*u*_(*θ* − Δ/2, *θ* + Δ/2) ≥ 4 (**Fig 6A**, iso-discriminability contours); we adopted a similar definition for the threshold in the masked condition, *T*_*m*_(*θ*). Comparison of the thresholds in the two conditions revealed that the mask elevated thresholds around the orientation of the mask (at 90 deg) (**Fig 6A**). Interestingly, the thresholds around the orientation orthogonal to the mask (0 deg) were not affected. A similar result is obtained if we perform a similar analysis based on *d*_*u*_(*θ*, *φ*) and *d*_*m*_(*θ*, *φ*) and assume the noise is uniform (data not shown).

**Fig 6.**
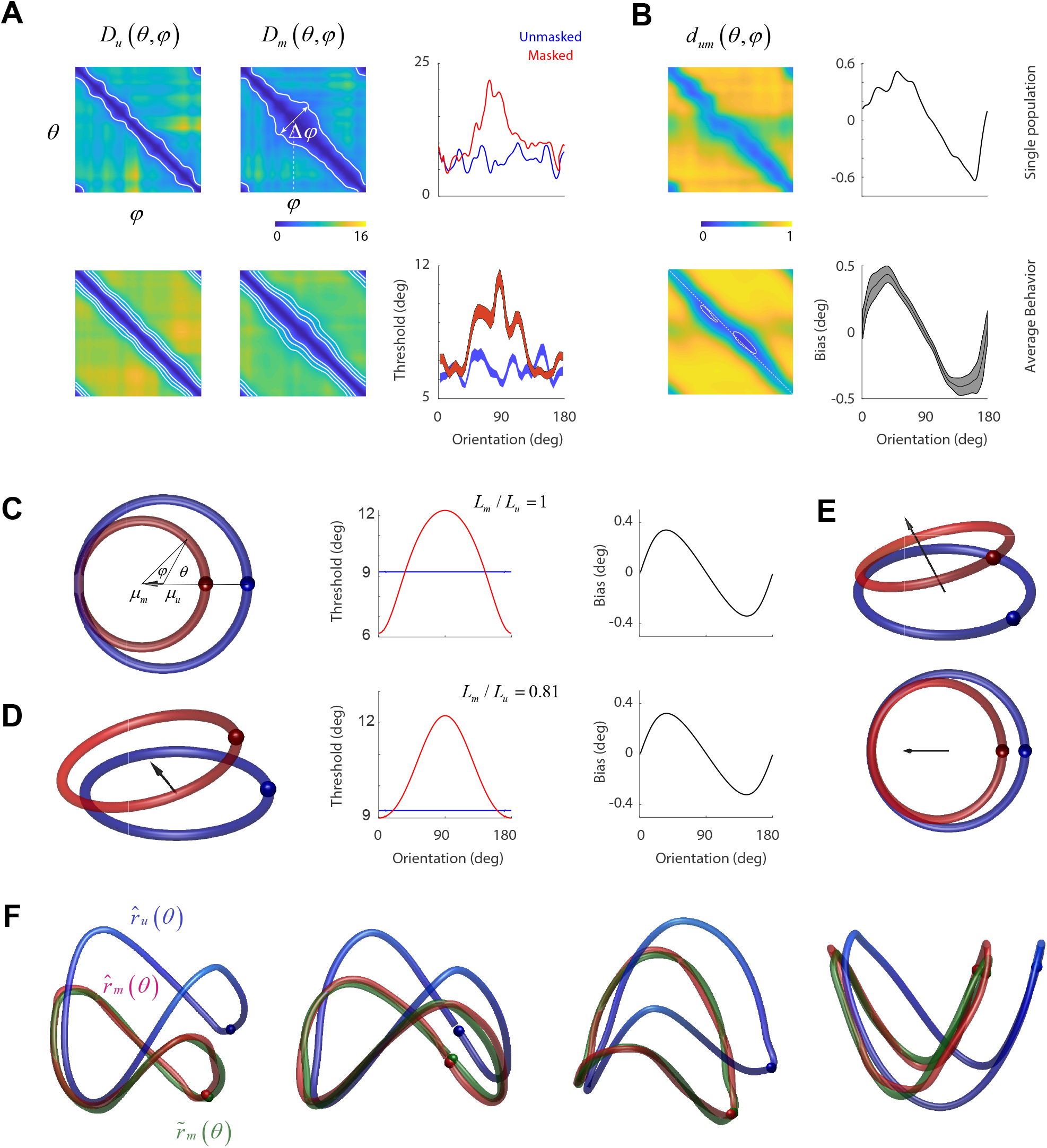
A geometric model of masking. **(A)** Discriminability (d-prime) between the representation of two orientations in unmasked (left panels) and masked (middle panels) conditions. The top panels show results for one experiment, while the ones at the bottom show the average across all our experiments. Iso-performance contour for the single experiment is shown at *d*′ = 4. The iso-performance contours for the average behavior is shown at levels of *d*′ = 4, 6, 8. The widening in the iso-performance contours in the masked condition reflect an increase in thresholds near the mask (which has an orientation of 90 deg). This is best shown in the panels on the right, which show the dependence of thresholds in masked (red) and unmasked (blue) conditions as a function of a base angle. In the average data the shaded areas represent ± 2 SEM. (**B**) Mutual distances and bias. Top panels show the mutual distance between orientations across masked and unmasked representations (*d*_*um*_) and the expected bias from a decoder based on the distances. The non-diagonal structure of *d*_*um*_ is more evident in the average data (bottom left panel), showing the locations of the minima of the main diagonal (white, dashed line). Bottom right panel shows the average bias across all our experiments. Shaded areas represent 2 SEM. (**C**) Two-dimensional geometric model of population coding^62^. The model assumes *r*_*u*_(*θ*) and *r*_*m*_(*θ*) are two circles in the plane. The displacement of their centers (white points) induce changes in the mutual distances inducing corresponding changes in threshold (middle panel) and bias (right panel). (**D**) The model can be extended by allowing displacement of the curves along a third dimension. (**E**) Two viewpoints of the same population activity in (**D**) but now normalized to yield 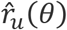 and 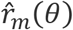. (**F**) Fits of an affine model to low dimensional representations of 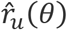 and 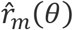 in four different experiments. In each case, 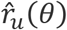 represents the population response in the unmasked condition (blue), 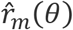 represents the population response in the masked condition (red), and 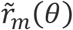 is the best fit to the response in the masked condition by means of an affine transform.

We then analyzed how the presence of the mask can lead to biases in estimates of orientation. We used a decoder based on population voting^57,61^. The estimated orientation was obtained as 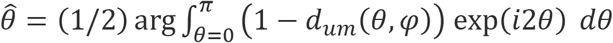. In other words, the population votes for each angle with a weight that depends on the distance to the representation of each angle in the unmasked orientation – the smaller the distance the strongest the vote. The bias is then 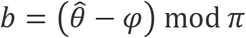. We observe that except at the orthogonal orientation the estimates are biased towards the orientation of the mask (**Fig 6B**). These biases arise because *d*_*um*_(*θ*, *φ*) does not have a non-diagonal structure -- the local minima of *d*_*um*_(*θ*, *φ*) occur slightly off the main diagonal (**Fig 6B**, bottom, white contours). Similar results are obtained using a simpler winner-takes-all decoder, where we pick 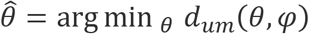.

### A geometric model for population transformations under masking

We tested if a simple geometric model^62^, originally developed to explain the effects of adaptation in psychophysical experiments, could explain our masking data (**Fig 6C**). The model assumes that in the unmasked condition the population response *r*_*u*_(*θ*) describes a trajectory around the unit circle and that the effect of the mask is to translate and scale this response to yield *r*_*m*_(*θ*). Translation is towards the population direction evoked by the mask, and the scaling is a typically a factor smaller than one. The model assumes that orientations are identified by the direction of the population vector, and that the decoder is unaware of the shift in the white point of the population between the two conditions. In other words, estimates of orientation are based on the direction of the population vector measured relative to the origin (which is also *μ*_*u*_ in this case) (**Fig 6C**). The model has only two parameters, the magnitude of the shift of the white point and a scaling factor. Its simplicity allows one to compute an analytical expression for both the threshold and the bias^62^ (see **Methods**). Indeed, this model captures some of the behavior of observed in the data. First, it reproduces the dependence of threshold with orientation in the masked condition, showing a maximum centered around the orientation of the mask. Second, it reproduces the shape of the bias reflecting an attraction towards the orientation of the mask.

The model, however, fails in three fundamental ways. First, in the model the population responses in both conditions lie within the same plane. As two independent vectors span the entire plane, it has to be the case that *r*_*m*_(*θ*) ∊ span{*r*_*u*_ (*θ*), *r*_*u*_(*π*/2)} (so long as *r*_*u*_(*θ*) ≠ *r*_*u*_(*π*/2)). In other words, this model satisfies linear combination^60^. However, we have already shown this is not the case in the data (**Fig 5D**). Second, both curves make a single revolution around the origin. This means that the lengths of the normalized responses are the same and equal to 2*π*, predicting a ratio *L*_*m*_/*L*_*u*_ = 1. Another way of stating this is that both 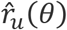 and 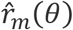 are unit circles, and that the two curves are different parametrizations of the same curve. However, the data show the ratios of the lengths to be significantly less than one (**Fig 5A**, tailed sign-test, *p* = 9.3 × 10^−10^). Third, the threshold is directly linked to how fast the population response changes its direction with angle, which is given by 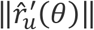 and 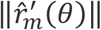. The faster the population direction rotates the lower the thresholds for discrimination. Faster rotation speeds However, as we just pointed out the average across all orientations is constant under this model, 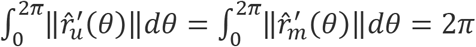. This means that if the mask increases discriminability for some orientations it must decrease it for others^58^. This is reflected in the fact that the threshold in the masked condition fluctuates around the mean for the unmasked condition (**Fig 6C**). The data, however, indicates that the effect of the mask was to impair the discriminability around the orientation of the mask, while there is little or no effect at the orthogonal orientation (**Fig 6A**, right column). The data refutes the prediction that increases in threshold at some orientations must be accompanied by decreases in threshold at other orientations (**Fig 6C**).

Can model be extended to account for our results? We know from the analysis of single cell responses that some neurons are unresponsive in the unmasked condition but respond robustly in the presence of a mask (**Fig 1B** and **2A**). This fact alone indicates the population responses in the masked and unmasked conditions do not lie within the same subspace. Thus, one way to extend the model is to allow the population responses to be displaced relative to each other along a third dimension (**Fig 6D**). Consider the response in the unmasked condition to be the unit circle and the one in the masked condition to be a the result of an affine transformation, *T*(*r*) = *αAr* + *t*, where *A* is an orthogonal matrix (representing a rotation), *α* is a scaling factor, and *t* a translation. It is then possible to find parameters of the transformation that reproduce the ratio between the lengths of the curves, as well as the dependence of discriminability and bias on orientation (**Fig 6D**, middle and right panels). An affine transformation can be represented in homogenous coordinates as *T*(*r*) = *Ar* where the population vector now has an extra dimension to allow for translation. We can then write the transformation of the *normalized* population responses as 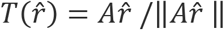, which we recognize as a projective linear transformation^63^ (**Fig 6E**). When the model is fit to the data in individual experiments, it nicely accounts for the observed transformations (**Fig 6F**).

## Discussion

Understanding how populations of neurons encode a physical attribute of a sensory stimulus, and how responses are transformed by contextual modulation is a central question of system neuroscience^49^. Here we considered the simpler question of how the orientation of a sinusoidal grating is transformed by an additive mask.

At the single cell level, we observed a wide range of responses (**Figs 1C,2**). Interestingly, we found a group of neurons that do not respond to gratings in the unmasked condition but respond strongly to plaids in the masked condition (**Fig 2A**). The maximal response of this type of these *plaid neurons*, occurs when the pattern is an orthogonal grating (**Fig 2B**). This finding implies that the responses in masked and unmasked conditions do not lie within the same subspace. This explains why the linear model (**Fig 5C**) and the 2D geometric model (**Fig 6C**) fail to account for the data. Grating and plaid cells are reminiscent of pattern and component cells^64,65^. We use different terms because the definitions are not equivalent. We note, however, that the pattern index used to classify cells as pattern/component correlates with the plaid/grating response we use here^66^ and that mouse primary visual cortex contains a larger proportion of pattern cells than found in non-human primates^67^. Thus, we suspect that the neurons engaged during masking that do not respond strongly to gratings in the masked condition could represent pattern cells.

We observed that plaid cells, when probed with a single component in the unmasked condition, responded optimally (albeit weakly) to the orientation of the mask (**Fig 2B**). While somewhat puzzling, the behavior in the masked condition might be explained if the addition of a grating orthogonal to a cell preferred orientation (as defined with single gratings) increases its response by releasing it from inhibition from oblique orientations in a ring model of orientation tuning^68^.

The main contribution of the present study is the introduction of a geometric approach to study contextual modulation of neural populations^48,62^. The analysis revealed that, despite a substantial heterogeneity in the behavior of individual cells, the map relating population responses in masked and unmasked conditions can be approximated as an affine transformation. When considering normalized responses, the corresponding map is a projective linear transformation^63^. The finding is so-far limited to masking, but we conjecture it may hold for other types of contextual modulation, such as interactions between the classical receptive field and the surround and sensory adaptation. Indeed, the a 2D model which accounts for psychophysical data on adaptation^62^ (**Fig 6C**) is an instance of an affine transform.

The geometric approach proved helpful in understanding several important properties of how population responses are modified by the introduction of a mask. First, it offered a straightforward test (and rejection) of a linear combination model^60^. The result could be easily understood as the mask moving the population activity out of its original subspace. Second, the analyses revealed that the transformation cannot be a reparameterization of the same curve, of which the 2D model is a special case^62^ (**Fig 6C**). The reason is that all reparameterizations leave the length of the curve invariant. In contrast, the mask was observed to shrink the length of the representation (**Fig 5A**). Third, we were able to show that the shift in the white point of the population is large compared to the radius of the curve (**Fig 5C**). This explains how a decoder which is unaware of such shift is bound to generate biased estimates. Finally, it clarified how a simple transformation can introduce changes in discriminability and bias in decoding (**Fig 6**).

Our finding of a white-point shift appears to be at odds with the idea that adaptation keeps the mean population response invariant (population homeostasis)^69^. In our terminology, population homeostasis would have predicted that *μ*_*u*_ = *μ*_*m*_, meaning no white-point shift. We suspect the reason for this discrepancy is rooted in the stimuli used. In the referenced study, a sequence of gratings with randomly chosen orientations was presented to the population. In one condition, the orientations were uniformly distributed; in the second condition, one orientation (the adapter) appeared more frequently than the others. In both conditions any one stimulus consists of a single grating. It is possible that such design failed to engage the plaid cells that clearly play an important role in shifting the white point. Similarly, a previous report^60^ selected cells to be analyzed only if their orientation tuning in response to a grating showed good selectivity (circular variance less than 0.85). Perhaps, plaid cells that were either unresponsive or weakly responsive to gratings failed to pass this criterion. The result would be biased towards gratings cells and it is possible that a linear combination model could be satisfactory when applied to this subpopulation of neurons.

Our findings indicate that analyzing the patterns of activity across large population of neurons we might be able to discover some general principles of sensory representation, including topological^70^ and geometrical structure, that are undetectable at the single cell level. These patterns can allow us to describe the transformations of representations in a simple way, as appeared to be the case for masking. Novel technologies that allow the recording of hundred or thousands of neurons simultaneously provide an ideal test bed for these ideas.

## Methods

### Animals

All procedures were approved by UCLA’s Office of Animal Research Oversight (the Institutional Animal Care and Use Committee) and in accord with guidelines set by the US National Institutes of Health. A total of 5 tetO-GCaMP6s mice (Jackson Labs), both male (3) and female (2), aged P35-56, were used in this study. Mice were housed in groups of 2-3, in reversed light cycle. Animals were naïve subjects with no prior history of participation in research studies. We imaged 30 different fields, and obtained data for 3920 cells, for a median of 111 cells per field (range: 50 to 275).

### Surgery

Carprofen and buprenorphine analgesia were administered pre-operatively. Mice were then anesthetized with isoflurane (4-5% induction; 1.5-2% surgery). Core body temperature was maintained at 37.5C using a feedback heating system. Eyes were coated with a thin layer of ophthalmic ointment to prevent desiccation. Anesthetized mice were mounted in a stereotaxic apparatus. Blunt ear bars were placed in the external auditory meatus to immobilize the head. A portion of the scalp overlying the two hemispheres of the cortex (approximately 8mm by 6mm) was removed to expose the underlying skull. After the skull was exposed it was dried and covered by a thin layer of Vetbond. After the Vetbond dried (approximately 15 min), it provided a stable and solid surface to affix the aluminum bracket with dental acrylic. The bracket was then affixed to the skull and the margins sealed with Vetbond and dental acrylic to prevent infections.

### Imaging and signal extraction

Imaging was performed using a resonant, two-photon microscope (Neurolabware, Los Angeles, CA) controlled by Scanbox acquisition software (Scanbox, Los Angeles, CA). The light source was a Coherent Chameleon Ultra II laser (Coherent Inc, Santa Clara, CA) running at 920nm. The objective was an x16 water immersion lens (Nikon, 0.8NA, 3mm working distance). The microscope frame rate was 15.6Hz (512 lines with a resonant mirror at 8kHz). We monitored locomotion using a rotary, optical encoder (US Digital, Vancouver, WA) connected to the rotation axel. The quadrature encoder was read by an Arduino board. We performed motion stabilization of the images, followed by signal extraction and de-convolution to estimate the spiking of neurons. The details of these methods are described elsewhere^51,55,71^. We used the average delay (387ms) measured in reverse correlation experiments to correct for the stimulus-response delay in the data^71^.

### Visual stimulation

We measured the responses of neural populations in mouse primary visual cortex using two-photon imaging in tetO-GCaMP6s mice (Jackson Labs #024742). The visual stimulus consisted of two conditions. In the first, unmasked condition, a sinusoidal grating (50% contrast and a spatial frequency in the range 0.04–0.06cpd) was presented with an orientation that changed linearly with time *θ* = *πt*/*T*, and a period *T* = 10s. The spatial phase of the grating was updated every *T*_*ϕ*_ = 783 msec by *ϕ* ← *ϕ* + *π*/2 + *n*, where *n* was a random variable distributed uniformly *n*~*U*(−*π*/8, +*π*/8). In other words, the grating underwent a “noisy contrast reversal” as its orientation changed continuously with time. This ensured that different spatial phases were present during different cycles of the stimulus. The unmasked condition was displayed for 15 min for a total of 90 cycles around the orientation domain. Immediately after, we added a vertical mask. The vertical mask also underwent a noisy contrast reversal with a period of 717 msec. A TTL pulse was generated by an Arduino board at the beginning of each stimulus cycle. The pulse was sampled by the microscope and time-stamped with the frame and line number being scanned at that time.

The screen was calibrated using a Photo-Research (Chatsworth, CA) PR-650 spectro-radiometer, and the result used to generate the appropriate gamma corrections for the red, green and blue components via an nVidia Quadro K4000 graphics card. The contrast of the stimulus was 99%. The center of the monitor was positioned with the center of the receptive field population for the eye contralateral to the cortical hemisphere under consideration. The locations of the receptive fields were estimated by an automated process where localized, flickering checkerboards patches, appeared at randomized locations within the screen. This measurement was performed at the beginning of each imaging session to ensure the centering of receptive fields on the monitor.

### Data analysis

We computed discriminability between two angles *θ* and *φ* as follows. Consider the responses in the unmasked condition. Let 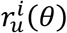 be the response of the population in the *i-th* cycle to a given orientation and let *μ*_*u*_(*θ*) be the mean population response across all trials. We define 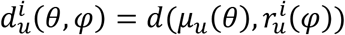. We then compute the indices 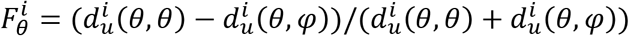 and 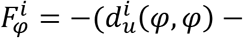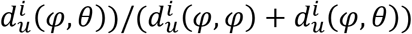. Finally, we compute *D*_*u*_(*θ*, *φ*) as the difference in the means of these distributions normalized by the average standard deviation. The same calculation was applied for the masked condition.

### Fitting the geometric model to experimental data

Note that the affine model in *d* dimensions has a total of *d*(*d* + 1) parameters. Our data consists of *s* = 155 equally spaced samples (10 sec period at 15.5 fps) of the continuous curves *r*_*u*_(*θ*) and *r*_*m*_(*θ*). Each sample provides *d* constraints on the transform. Thus, we must have *d*(*d* + 1) ≤ *ds* or (*d* + 1) ≤ *s* to ensure the problem is not under-constrained. We handled this constraint by first projecting the data onto into *R*^3^ using the first three components in the SVD factorization of the data and subsequently fit the lower-dimensional embedding of the curves (**Fig 6F**). Note that projection is a linear operation. Thus, if the data conformed to an affine model in the high-dimensional space, it should also do it its low dimensional embedding (no matter how much distortion we are imposing by the projection). This can be easily shown using a basis set corresponding to the canonical form of the projection. The reverse, of course, it not necessarily true.

### Analytic computation of threshold and bias

In the simple geometric model of **Fig 6C** it is possible to compute the threshold and bias. Consider a two-dimensional population code for orientation in the unmasked condition *r*_*u*_(*θ*) = (cos *θ*, sin *θ*), which is transformed by a scaling and translation along the x-axis under the presence of the mask *r*_*m*_(*θ*) = (*a* + *b* cos *θ*, *b* sin *θ*). Then, the velocity of *r*_*m*_(*θ*) is

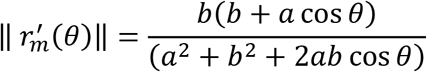

The threshold will be inversely proportional to the velocity 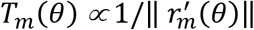. Given a population direction in the masked condition, which in the plane is simply given by an angle *φ*, a decoder without knowledge of the white point shift will estimate the orientation by measuring the angle *θ* formed between the population vector with respect to *μ*_*u*_ (**Fig 6C**), which a little geometry shows it is given by *θ* = arctan((*a* + *b* cos *φ*)/ sin *φ*). Thus, the bias is given by

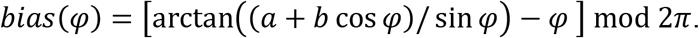

## Acknowledgements

I thank Elaine Tring for help in surgical preparation and data collection.

